# Confusions regarding stochastic fluctuations and accumulators in spontaneous movements

**DOI:** 10.1101/2021.06.04.447111

**Authors:** Carsten Bogler, Bojana Grujičić, John-Dylan Haynes

## Abstract

Experiments on choice-predictive brain signals have played an important role in the debate on free will. In a seminal study, Benjamin Libet and colleagues found that a negative-going EEG signal, the readiness potential (RP), can be observed over motor-related brain regions hundreds of ms before the retrospectively reported time of the conscious decision to move. If the onset of the readiness potential is taken as an indicator of the “brain’s decision to move” this could mean that this decision to move is made early, by unconscious brain activity, rather than later, at the time when the subject believes to have decided. However, an alternative kind of interpretation, involving ongoing stochastic fluctuations, has recently been brought to light. One such model, the stochastic decision model (SDM), takes its inspiration from accumulator models of perceptual decision making. It suggests that the RP originates from an accumulation of ongoing stochastic fluctuations. In this view the decision happens only at a much later stage when an accumulated noisy signal (plus imperative) reaches a threshold. Here we clarify a number of confusions regarding both the evidence for the stochastic decision model as well as the interpretation that it offers. We will explore several points that we feel are in need of clarification: **(a)** that the empirical evidence for the role of stochastic fluctuations is so far only indirect; **(b)** that the relevance of evidence from animal studies is unclear; **(c)** that a model that is deterministic during the accumulation stage can explain the data in a similar way; **(d)** that the primary focus in the literature has been on the role of random fluctuations whereas the deterministic aspects of the model have been largely ignored; **(e)** that contrary to the original interpretation the deterministic component of the model is the dominant input into the accumulator; **(f)** that there is confusion regarding the role of “imperative” and “evidence” in the SDM and its link to perceptual decision making; and finally **(g)** as with other stochastic accumulator processes the question of whether the decision happens early or late depends on the nature of the noise fluctuations, specifically, whether they reflect “absolute” or “epistemic” randomness. Our aim is not to rehabilitate the role of the RP in the free will debate. Rather we aim to address some confusions regarding the evidence for accumulators playing a role in these preparatory brain processes.

## Introduction

Throughout the day we have to make a multitude of decisions about *external* stimuli. For example, when we see a car crossing our lane on the highway, we step on the break to avoid a collision. An important factor is the level or quality of sensory information. For example, when driving in broad daylight we instantly see the dangerous car. But when it is foggy, we might be uncertain about whether it is a car or just a random pattern in the mist. In that case we might need to look at the pattern for a bit longer and gather evidence across time. A popular approach for explaining perceptual decision making (PDM) under such varying levels of sensory evidence is the accumulator model (Smith & Ratcliff, 2004). It formulates a mechanism that accumulates sensory evidence across time and thus gradually improves the accuracy of a sensory decision. When the buildup of evidence crosses a set threshold the decision is reached, and a reaction can be triggered. Most accumulator models involve two key variables that are combined in an additive fashion: the first term is the *sensory evidence* in each time step that reflects a single sample of information about the external stimulus; the second term is *internal noise* that accounts for variability in responses. The accumulator adds both terms, the evidence and the noise, as inputs to its ongoing total evidence tally. So, both the external evidence and the internal noise contribute to the decision. When the external information is high (as in broad daylight) the process is dominated by the evidence, and the threshold can be reached quickly. When the external information is low (as in fog) the process is dominated by the internal noise, and it takes longer to reach a decision. The model may also include a leak term so that the total evidence slowly decays if it is not refreshed (Usher & McClelland, 2001).

In recent years, this approach has also been used to explain the neural mechanisms underlying simple, spontaneous voluntary actions (Schurger, 2018; Schurger et al., 2012). These movement decisions have been met with considerable interest in debates about free-will and volition (Libet, 1985). This is because such spontaneous decisions are preceded by a slow negative-going EEG signal, the so-called readiness potential (RP) (Kornhuber & Deecke, 1965) that appears to occur even before the time at which a person reports to have made a conscious decision to move (Libet et al., 1983). To give a very rough summary, a debate has centered on the following notion: if the brain “knows” that a decision will occur before a participant has consciously made up their mind, then this might mean that the decision has happened before the conscious mind became involved, which has been debated as a potential challenge to conscious free will (for discussions see Brass et al., 2019; Libet, 1985; Schurger et al., 2012, 2021). Here, we will not be interested in the free will debate, but in the mechanisms that occur before a self-initiated movement. According to the original interpretation, the onset of the RP is a “post-decisional” signal, meaning that the buildup begins once the decision to move has been made by the brain. In that view the early onset of the RP reflects an early decision of the brain that happens before consciousness kicks in (Schurger et al., 2021).

Recently, an alternative kind of explanation has been proposed that is based on ongoing slow random fluctuations in brain activity and that places the decision at a much later time. As we will see, a key difference here is that the RP, rather than being post-decisional, reflects a pre-decisional stage where the decision has not yet been made and during which random fluctuations play a role in determining the precise time at which the threshold is reached. One such model, the stochastic decision model (SDM) proposed by Schurger et al. (2012), takes its inspiration from the leaky accumulator model that is well-established in perceptual decision making (Usher & McClelland, 2001). However, now the noise term from the accumulator takes center stage. The idea is that the decision is determined largely by the accumulation of random internal fluctuations. Fluctuation-based accounts have long been used to explain Libet’s findings (Eccles, 1985; Libet, 1985; Ringo, 1985; Stamm, 1985). These older accounts (some of which are dualist) do not explicitly employ accumulators, but slowly fluctuating signals (note: a leaky accumulator is similar in some ways to a low-pass filter and generates similar slow fluctuations when provided with noisy input (Eliasmith & Anderson, 2004).

In the original version of the SDM (Schurger et al., 2012), two different sources provide input to a leaky accumulator. The first input stems from noise fluctuations, but these alone would not drive the signal over the decision bound within the typical time taken by participants to make a decision (i.e. model would not account for the distribution of waiting times). So, a second input is used that brings the process into the operating range close to the decision bound so that the accumulated internal fluctuations can spuriously drive the signal across the boundary. Or in their original words:

> *“In our model this solution amounts to simply shifting premotor activation up closer to the threshold for initiation of the instructed movement and waiting for a random threshold-crossing event.”* (Schurger et al., 2012, p. E2905).

In this version of the SDM, this process that drives the signal closer to the threshold is a constant input called an “urgency” or “imperative” signal, as it reflects the demand or imperative to move (Schurger et al., 2012). Depending on where the threshold is set, it is necessary because it prevents having to wait for a very long time for the decision (see below). Interestingly, this imperative signal is mathematically equivalent to the “evidence” signal in perceptual decision making, but it has a very different interpretation (see also below). Note that the word “urgency” used in Schurger et al. (2012) may have been confusing because of the different way that same word is sometimes used in the perceptual decision-making literature (Cisek et al., 2009). We thus follow Schurger (2018) and use the term “imperative” here.

In the following our primary aim is to clarify several points regarding the SDM that have led to confusions in the literature. While many authors correctly cite and discuss the architecture and the implications of the model, there still seem to be a lot of misunderstandings regarding several aspects. Our interest here is not to fully review the literature on the readiness potential, to re-introduce the readiness potential into the debate on free will, or to rule out the accumulator model as a potential mechanism for spontaneous actions. We will focus on the discussion of the mechanisms involved in movement initiation rather than the question of subjective experiences of volition. Our prime aim is to delineate more clearly what the SDM-related findings mean and what they don’t mean. We will provide some equations, but this paper should be approachable even without an in-depth understanding of the mathematical foundations. Readers who do not want to dive into the mathematical basics can jump over the next section.

### How does the accumulator model work?

In this section we will go into more detail about how the model works and which evidence is provided in support of it. We will focus on two papers, Schurger et al., (2012) and Schurger (2018), because these contain explicit mathematical formulations. They are both similar variants of the larger family of accumulator models (e.g. Smith & Ratcliff, 2004). We will reorder some aspects for easier readability, and we will use difference equations rather than their differential variants because all their modelling is done in discrete time steps *i*.

The key variable is the accumulated signal x_i_ (Fig. 1, left), often referred to as the “decision variable” in perceptual decision making. At the beginning of the trial this variable starts at x_0_=0 (a starting bias is not used) in their implementation. On every time step an increment or decrement Δ*x*_*i*_ is added to *x*_*i*_ and when *x*_*i*_ reaches a threshold *β* a movement is triggered at time point T also referred to as the waiting time. (Please note: In a realistic brain there is still a delay between the time T when the movement command is sent into the motor system, say down the spinal cord, and the time when the movement begins in the muscles, see Schurger et al., 2012).

**Fig. 1:**
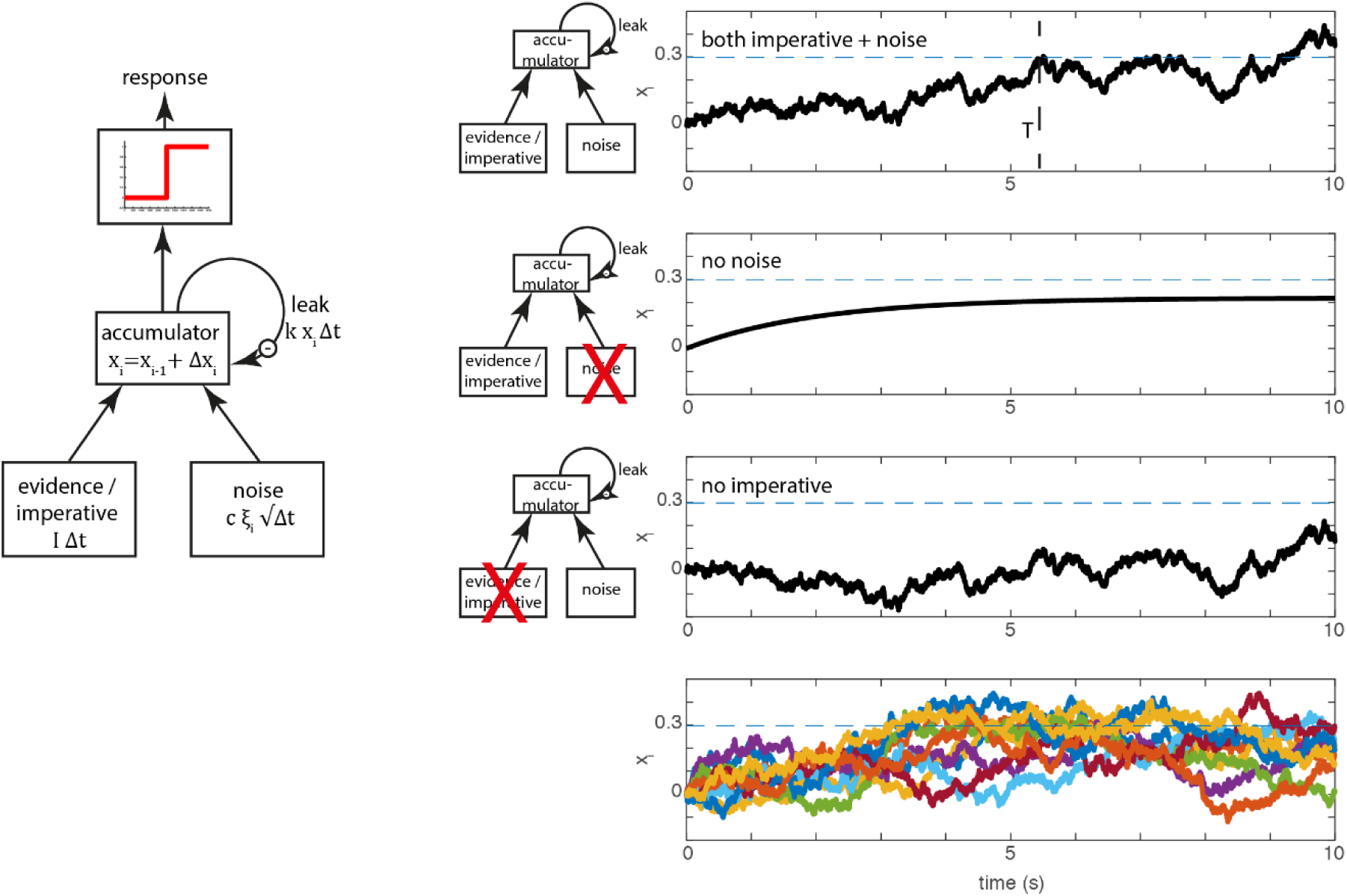
Basic accumulator model. **Left:** In perceptual decision making at each time step, two variables, evidence and noise are added to a leaky accumulator. When the output of the accumulator reaches a certain threshold (red) a report is triggered. In spontaneous movement, the evidence is replaced by an imperative term that can help bringing the signal closer to the threshold. **Right:** Examples of the stochastic decision model (SDM) (Schurger et al., 2012). The first three rows on the right show the behavior of the model in a single trial, separately for the full model (top), only the evidence/imperative and leak with noise removed (second row) and only the noise with leak (third row). The dashed horizontal line is the threshold *β*. The bottom row shows 10 trials, which clearly highlights the variability in individual trial accumulator trajectories. Note that in this original model, the imperative with leak will not drive the accumulator beyond the threshold, and the noise with leak will take an implausibly long time to drive the accumulator over the bound. Both terms together bring the signal across the threshold (at T, top row), which then triggers a movement with a distribution of reaction times that matches the waiting times of the participants until they press the button.

At each time step *x*_*i*_ is updated by Δ*x*_*i*_ based on the following equation (rewritten from the original in a slightly more explicit form):

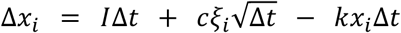

This means that the increment/decrement that is added to *x*_*i*_ on each trial depends on three additive components:

1. The first term, *I*Δ*t*, is a constant that is referred to as the “imperative” in the SDM. This is mathematically equivalent to the (mean) evidence in accumulator models of perceptual decision making, but it has a somewhat different interpretation.
2. The second term, 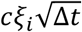, reflects internal Gaussian noise *ξ*_*i*_that is scaled by *c* and 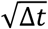 (in the SDM both Δ*t* and *c* are fixed scaling constants).
3. The third term, *kx*_*i*_Δ*t*, represents leakage, with the leak constant *k* scaled with another constant Δ*t*. Thus, the accumulated signal is reduced by a constant proportion of *x*_*i*_ on each time step.

When the accumulated signal *x*_*i*_ crosses the threshold *β*, a motor command is triggered. Figure 1 (left) shows the operation of the model expressed as a more conventional box and arrows model. There, the three inputs from above are shown as arrows feeding into the accumulator. Let’s look at the behavior of a single trial (Fig. 1, top right; Fig. 1, bottom right, shows this process for a large number of trials). In every trial the accumulation starts at the first step at *x*_0_ = 0 (other accumulator models sometimes introduce a starting bias here). At every time step the increment (or decrement) Δ*x*_*i*_ is added to *x*_*i*_, resulting in a noisy drift towards the decision boundary. At some point the accumulated signal crosses the decision boundary (Fig. 1, top right, dashed line) and triggers a response at latency T.

The next two rows of the figure show the consequences of removing the imperative (i.e. the constant) term versus the noise term. If the noise term is removed the signal *x*_*i*_ rises and depending on the parameters of the model asymptotes below the threshold (as shown here, with the parameters in Schurger et al., 2012) or it crosses the threshold (Schurger, 2018). If the imperative term is removed and the other variables are kept the same, the signal meanders around for a long time and at some point crosses the threshold, but with an implausibly long latency. Thus, in this variant of the model, both the imperative and the noise are involved in bringing the system to the threshold.

### Differences between the spontaneous motor decision model and perceptual decision making

The SDM for endogenous decisions is *mathematically* equivalent to a variant of the perceptual accumulator model (Usher & McClelland, 2001). The two scenarios differ only regarding the interpretation of the parameter *I*. In perceptual decisions, the drift term *I* refers to the mean sensory evidence. Normally, in perceptual decision making, the constant term, the sensory evidence, is the main driving factor towards the decision boundary. The noise component is sometimes referred to as reflecting moment-to-moment *changes* in evidence, but on average and by itself the noise component does not contribute any evidence at all because it is mean-centered. In perceptual decision making, when the evidence is 0, the behavior of the accumulator is governed by the noise term (see Fig. 1, right).

We would like to highlight two points of the accumulator model in perceptual decision making. First, in the model as formulated here, for a given evidence level the drift *I* is a constant (see e.g. Ratcliff, 1978, for variations on this assumption). It reflects the mean amount of sensory evidence that some neurons are encoding about an external stimulus property. For example, a motion stimulus of a specific coherency level on an external monitor will lead to a constant representation of momentary evidence about this motion direction in motion area MT in the brain, and this evidence is summed up by the accumulator. Second, the term “evidence” here means that the signal in MT has information *about another property*, the external motion stimulus. This evidence can also be very small or even 0 in case of very weak or no sensory evidence.

So how does this perceptual decision-making model transfer to spontaneous movements? In a review paper Schurger and colleagues state:

> “*A strength of SDMs [stochastic decision models] is that they provide a unifying story that seamlessly allows agents to move between reason-driven and random decisions, as the spontaneous action case is <u>just an SDM driven by noise</u> in the absence of evidence/reasons.*” (Schurger et al., 2021, p. 10, underline added).

Based on this statement one might think that the spontaneous movement model (SDM) is based on the perceptual decision-making model, but with zero evidence, with only the noise active, thus making it similar to perceptual guessing. In line with this, we have shown previously that perceptual guesses (perceptual decisions with no sensory evidence) and spontaneous decisions indeed elicit similar activation patterns in posterior parietal cortex (Bode et al., 2013). It is also a reasonable hypothesis to explain free choices using an accumulator that is provided with only stochastic input but no evidence. However, in the accumulator model of Schurger et al. (2012) the constant drift/evidence term *I* was explicitly not set to zero, but was rededicated and given a new role as an “imperative” parameter. It was subsequently confirmed by the authors that the accumulation of sub-threshold noise alone would not be sufficient to fit the behavioral and brain data (see Fig. 2) (Guevara Erra et al., 2019). The roles of imperative and noise in the model have been implied to reflect a two-stage sequential process:

> *“*In our model this solution amounts to simply shifting premotor activation up closer to the threshold for initiation of the instructed movement and waiting for a random threshold-crossing event.*”* (Schurger et al., 2012, p. E2905).

**Fig. 2:**
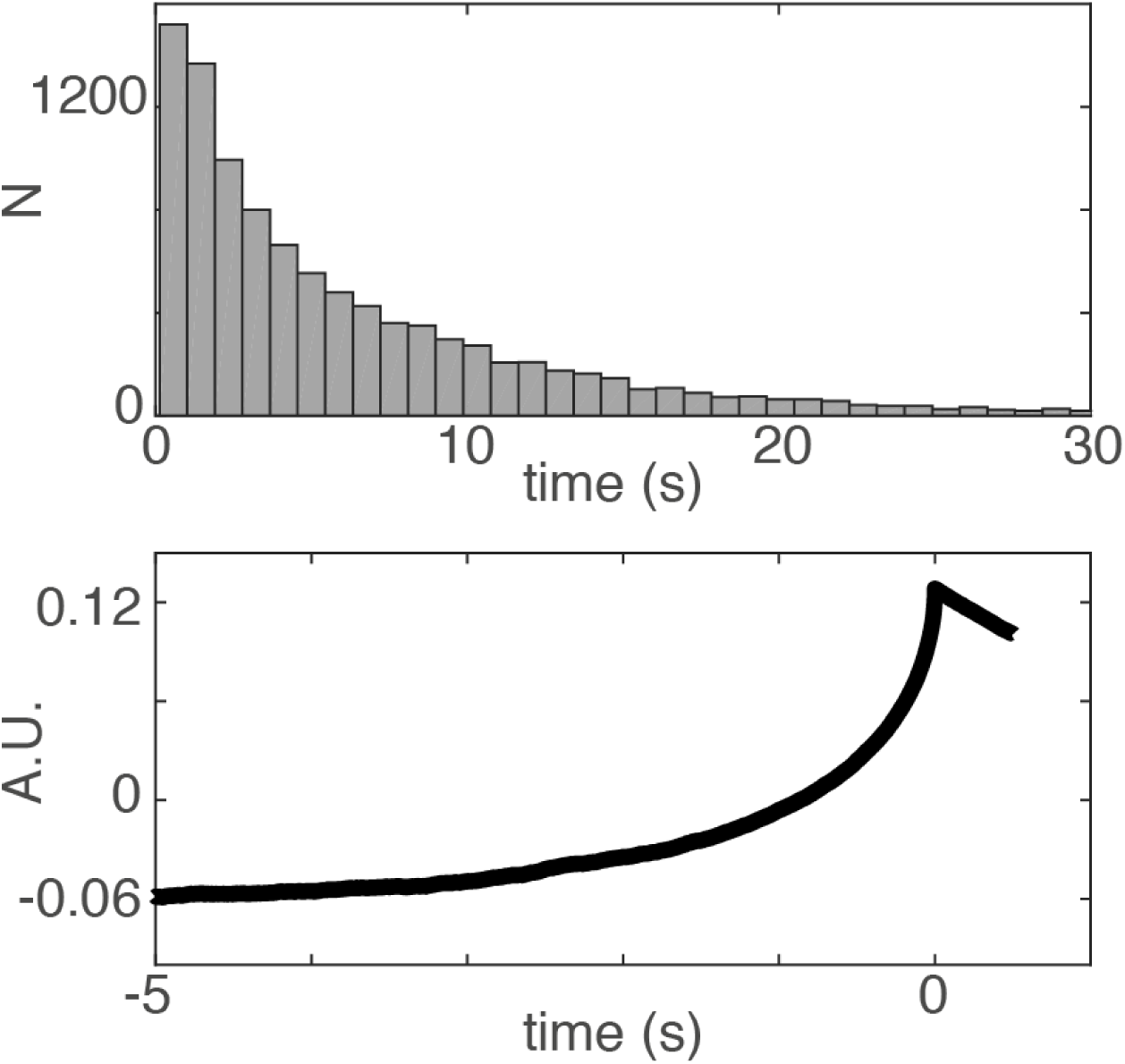
The distribution of waiting times (**top**) in a variant of the Schurger et al. (2012) model where the imperative was set to 0 and the other parameters were adjusted (threshold was adapted to 0.1265 corresponding to the 80^th^ percentile of the output amplitude, Schurger et al., 2012). With this adjustment, the distribution of waiting times does not match the shape of the empirical waiting times of subjects performing the task. However, the RP (**bottom**) from that model has typical RP characteristics. Thus the imperative is important in order to fit the behavioral data (see also Guevara Erra et al., 2019). But even with zero imperative the SDM can explain the typical shape of the RP.

We will see more examples of this notion below and demonstrate that both claims may be misinterpreted: the contribution of the imperative is substantial (see below), and there are no two disjunct stages, pre-stationary and stationary, in the process described by the model. Importantly, half of the decisions happen during the early stage where the imperative effect is moving the signal closer to the threshold (see below and Fig. 9). Please note that an imperative signal is only one way to bring the accumulated signal and the threshold closer together. Alternatives could potentially be changing the starting point of the accumulation process or the decision boundary itself (but see Guevara Erra et al., 2019) or progressively “collapsing” the decision boundary across time (Hawkins et al., 2015). Thus, it might be better to speak of a two-component rather than a two-stage process.

### How is the accumulator linked to the readiness potential?

In order to provide support for the model, Schurger et al., (2012) show that it provides a potential explanation of the readiness potential. The idea is that the RP emerges from averaging the trajectory of the accumulated signal *x*_*i*_ *backwards* from when it reaches the threshold (Fig. 3). Importantly, all the fitting here is done based on the *average* RP, i.e. by averaging across many trials (for single-trial extraction of RPs see e.g. Schultze-Kraft et al., 2016). Fig. 3 shows this principle and plots some sample trajectories using the best fitting parameters from Schurger et al. (2012). These are obtained by fitting the waiting time distribution predicted by the model to the empirical waiting time distribution observed in the behavioral data. Please note that the waiting time distributions are subject to additional transformations before being compared (Schurger et al., 2012, p. E2906). Given those specific parameters the model also predicts the readiness potential. Also note that the readiness potential only reflects the final stage of the modelled decision-making process. Thus, a distinction has to be made about the claims made by the entire decision-making model and the claims made relating to predicting the shape of the readiness potential.

**Fig. 3:**
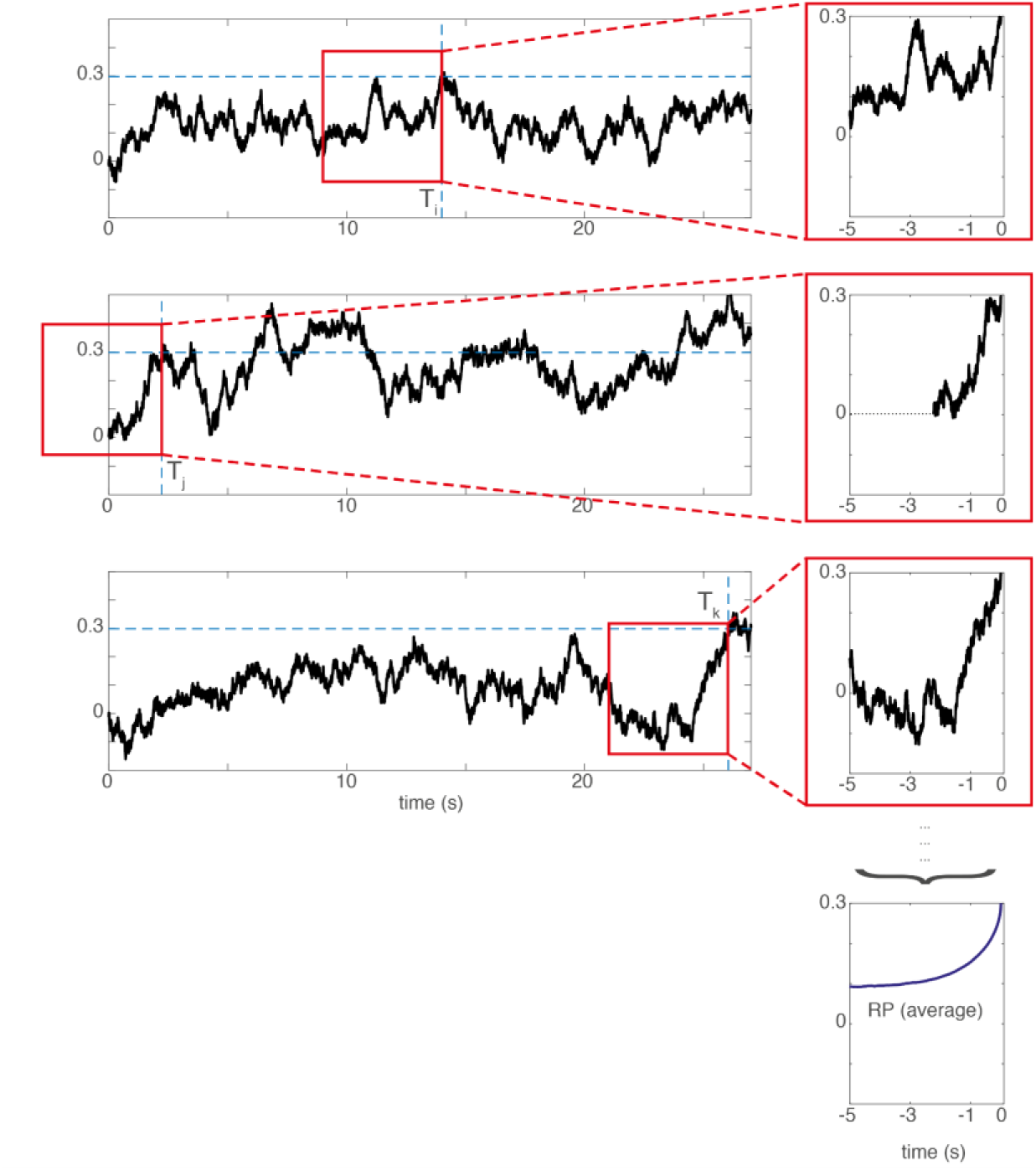
The SDM assumes that the readiness potential reflects individual trajectories of the accumulated signal *backwards*-averaged from the time of threshold crossing (T). The left shows artificially generated trajectories of a hypothetical accumulator signal *x*_*i*_ in three different trials. The red box shows the 5 second averaging time window averaged backwards from the threshold crossing that can be seen on the rightmost border. The top three panels on the right show the signal in the red window enlarged and temporally aligned. The bottom right panel shows the average of these threshold-crossing-aligned trajectories across 1000 trials. This curve can fit the shape of a readiness potential. When the threshold crossing happens early in the trial (2nd row), the missing values are left out in the average (i.e. they are coded in Matlab as NaN, “not a number”). Please note that the signal trajectories are latent variables of the model, but they are not measured directly. The model fit is conducted at the level of the RP averaged across 1000 trials. Also, see (Schurger, 2018) for different assumptions underlying the spectral nature of these noise fluctuations and for different architectures of the model. Please note that the RP directly derived from the model is positive-going because the threshold is positive as in the original paper. There the time course is sign-reversed to match the empirical RP, which is a negative-going voltage deflection.

### Is there direct empirical evidence for the role of noise fluctuations in the RP?

Next we would like to address a number of confusions that seem to have originated in the literature on the SDM. For example, summaries provided in various papers give the impression that the empirical analysis of the EEG data and of behavioral waiting times provides *direct* evidence for the involvement of fluctuations in the decision process (see below). This is not the case. Instead, the fluctuations constitute a latent variable of the model that is not directly measured, as is the case in many neuroimaging studies that employ unobserved variables for modelling. The fluctuating time courses were not measured at the single-trial level. They are hypothetical and are used as part of a model to predict average properties of the recorded data. The gap in resolution between model variables (e.g. noise time courses of hypothetical units) and measured variables (EEG signals) is not unusual in modelling of neuroimaging data. For example dynamic causal modelling (DCM) (Friston et al., 2003) also models measured data using a number of latent variables at finer levels of temporal resolution, however this is always done at the level of raw time series data, not on averages.

To date there is no directly-measured evidence for a role of stochastic fluctuations in generating the readiness potential. Thus, the question of whether arbitrary free choices (like those made in the Libet paradigm), and the accompanying RP, do in fact involve ongoing stochastic fluctuations is not yet settled. Furthermore, in recent years there have been substantial challenges to the ubiquity and nature of a core feature of the SDM, that is the accumulation process: Especially in situations where sensory evidence is brief rather than distributed across time, accumulation might not always take place (Thorpe et al., 1996; Uchida & Mainen, 2003), in other than in trivial ways (obviously one could debate whether e.g. the superposition of postsynaptic potentials constitutes “accumulation”). Furthermore, the true dynamics of information processing during decision-making might be difficult to infer from average data (e.g. Latimer et al., 2015). One solution might be to obtain invasive recordings in human patients, insofar as possible (see below), as in Fried et al. (2011)(see below).

### Is the model supported by invasive recordings in animals?

In order to provide more direct evidence, the authors point to converging studies on animals. Potential evidence for the neural implementation of a SDM for endogenous tasks was reported by Murakami et al. (2014). That study investigates spontaneous movements in an intertemporal choice task in rats. After a go-signal, rats are given a choice between an immediate water reward or, if they wait for a delayed second signal, a much higher reward. Sometimes rats wait a bit, but then spontaneously abort and go for the smaller immediate reward. These choices on “impatient trials” are considered endogenous because there is no immediate trigger to move. For these trials they make two observations: (a) The activity in one selectively chosen population of neurons (P1) in rat motor area M2 rises sharply in the last few hundred ms before the movement. They interpret this as an accumulated evidence signal; (b) In a separate selected set of neurons (P2), some neurons (P3⊆P2) show activity that is predictive of the waiting time early in the trial.

At first sight the similarities could be seen as providing support for the SDM. A careful look, however, shows that the superficial appearance of similarity may be misleading. By inspection of Fig. 5c,d in (Murakami et al., 2014) one can see that a *majority of predictive time periods are around the start of the trial or even before the onset of the trial*, which is the opposite of what would be expected in the case of an accumulator model. We will return to this later. Furthermore, the signal is transient. Thus, this specific signal does not seem to map on to any variable of the SDM model, neither to a continuous stochastic or constant input, nor to an accumulator continuously integrating input across the trial. Instead, it could simply reflect a cognitively interpretable bias signal, such as an expectation on that trial of when the delayed reward will occur. If anything, a different subpopulation of neurons exhibits a ramping-like behavior, but mostly towards the end of the trial. However, there is a considerable temporal dissociation between the time where most of the time windows are informative, and the time when the putative accumulator in their data ramps towards threshold (compare Murakami et al. 2014, Figs. 4 & 5). If the predictive signal feeds into the accumulation one would expect it to appear close in time to the steepest increase in the accumulated signal. In their decision model, the signal from each contributing neuron is only collected in a single brief time window, and otherwise ignored (see their p. 1584). Thus, the accumulator model proposed by Murakami is quite different from previous models.

**Figure 4:**
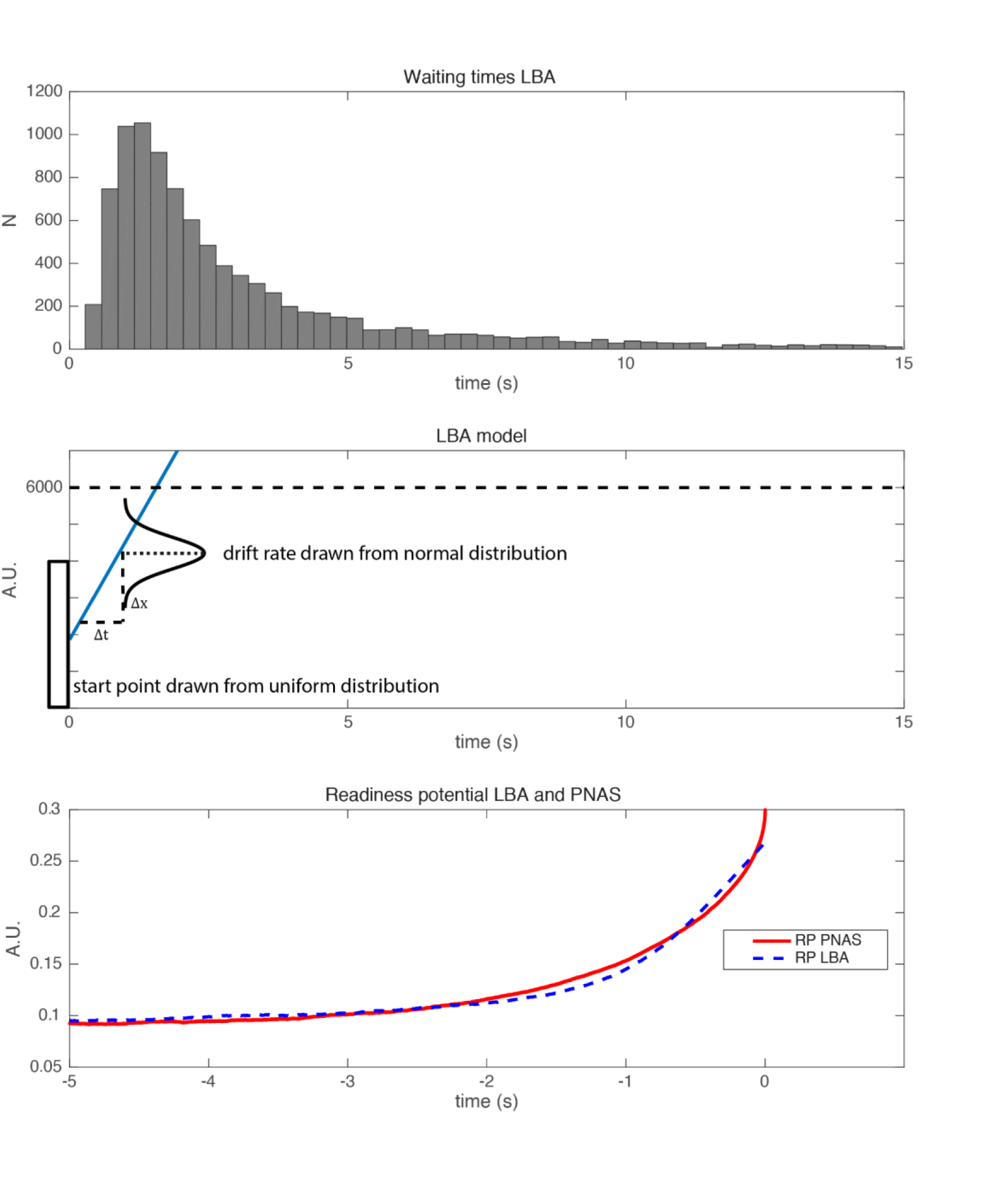
Simulation of a linear ballistic accumulator model (LBA). **Top:** Waiting times generated with the LBA. **Middle:** Schematic plot of the linear ballistic accumulator model. The starting position is drawn from a uniform distribution (between 0 and 4000). The drift rate is drawn from a normal distribution (mean = 1, std = 2). The threshold is at 6000. **Bottom:** Readiness potentials generated with the LBA (blue) and the original SDM (red). Please note that the RP of the LBA is scaled in order to match the RP of the PNAS model (Schurger et al., 2012). Please note that in order to match empirical RPs both models involve additional scaling factors, which also ensure that the polarity of the time course is inverted to match the polarity of the empirical RP.

**Figure 5.**
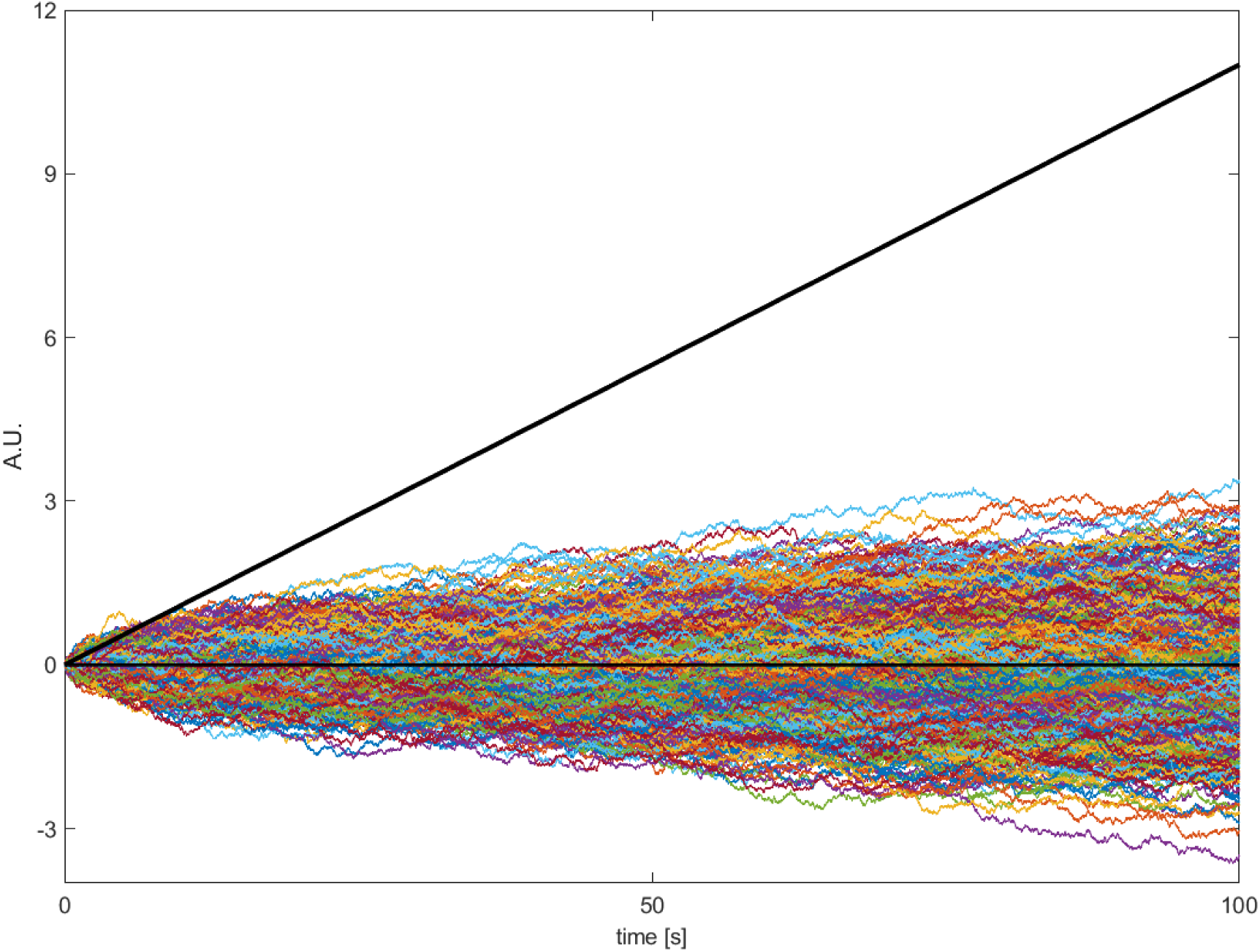
The cumulative input into the accumulator across the trial, plotted separately for stochastic and constant input. The cumulated imperative (∑ *I*Δ*t*) is deterministic for every trial (bold black line) and thus there is only one trace. The cumulated noise (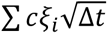, colored lines) is shown here for 1000 different simulated trials. Please note that the cumulated noise across simulated trials is on average zero and the amplitude of single trials can be positive and negative. The noise is negatively correlated with the waiting time. Positive values will push the total signal over the threshold quicker whereas negative values will work against reaching the threshold and lead to longer waiting times. Most importantly, it can be easily seen that the cumulated imperative is almost always higher compared to the cumulated noise, especially for later time points.

### Are random fluctuations necessary in order to account for the data?

Given that the fluctuations have not been directly measured, but only indirectly inferred, it would be interesting to know whether a simpler model, potentially without fluctuations during the accumulation process, could in principle also explain the RP. Of particular interest would be a model that is compatible with the early, choice-predictive signals observed in the study by Murakami et al. (2014) mentioned in the previous section. There is indeed one such model, the linear ballistic accumulator (LBA), an accumulator model that was originally developed for perceptual decision making (Brown & Heathcote, 2008). In contrast, this does not have stochastic fluctuations during the trial, but replaces those with a randomly selected drift rate and starting bias that are only determined in a single step at the beginning of each trial. The LBA (Fig. 4, middle) is a simplified version of the standard accumulator model. The difference is that the drift rate *I*, as well as a starting bias, is drawn from a random distribution only once at the beginning of each trial. Thus, it is not subject to noise fluctuations within the rest of a trial at all. The drift rate varies across trials, which could reflect e.g. differences in attention (in the case of perceptual decision making) or differences in motivation or impulsivity (in the case of spontaneous movements). Interestingly, this fluctuation-free model makes very similar predictions for features of perceptual decisions to the accumulator. Importantly, it predicts the typical heavy-tailed reaction time / waiting time distribution. When used in a similar way to predict readiness potentials, the ballistic accumulator model also provides a good fit to the RP, despite its simplicity and the absence of random fluctuations during the trial (Fig. 4). Please note that for this simple LBA model the whole process is pre-determined once the trial is started. Previously there has been the idea that at the beginning of the trial there is a *general* decision to move but the exact time is left open, and then subsequently during the trial there is a “decision to move now” that is the final commitment to immediately move (Schurger et al., 2012). The LBA would constitute a different view where some kind of early (or potentially unconscious) decision is made at the beginning of the trial to move at time T in the future.

In fact, one could question whether the LBA really is an accumulator at all, despite providing a good fit to the data and carrying the label “accumulator” in its name. The answer to “when the decision is actually made” is very different for the LBA than for the SDM, because the key factor that determines the outcome is present already at the beginning of the trial. As mentioned above, this model is somewhat more compatible with the data by Murakami et al. (2014) because it predicts that the response time is encoded in brain signals already at the beginning of the trial. Thus, in the absence of direct tests of the link between fluctuations and RPs, both early and late decision models appear to be equally plausible (for further discussion see below).

Please note that the LBA model makes an important similar prediction to the SDM. When participants were interrupted by a click in the waiting period then the response to that click was faster when the EEG signal was more negative (Schurger et al., 2012). This is also predicted by the LBA. The more the signal has approached the threshold the shorter a motor reaction time would be if the accumulation processes for endogenous and exogenous movements share this common path.

### What are the relative contributions of “noise” and “imperative”?

In this section we will try to clarify the relative contributions of noise and imperative signals in the accumulation process. Note that Schurger et al. (2012) and many subsequent summaries frequently primarily focus on the fluctuations and largely ignore the constant component. While they are indeed mentioned in the original manuscript, there are also other statements that focus solely on random fluctuations:

> *“One simple solution, given these instructions, is to apply the same accumulator-plus-threshold decision mechanism, but fed <u>solely</u> with internal physiological noise.”* (Schurger et al., 2012, p. E2905; underline added).

In a subsequent paper they say:

> “[…] *when actions are initiated spontaneously rather than in response to a sensory cue, the process of integration to bound is <u>dominated by</u> ongoing stochastic fluctuations in neural activity […]”* (Schurger et al., 2016, p. 78, underline added).
>
> “*In the case of spontaneous self-initiated movement there is no sensory evidence, so the process is <u>dominated by</u> internal noise.*” (Schurger et al., 2016, p. 77, underline added).

And even later as we have seen above:

> *“[…] the spontaneous action case is just an SDM <u>driven by noise in the absence of evidence/reasons</u>.”* (Schurger et al., 2021, p. 10, underline added).

Subsequently, many summaries of the findings ignore the role of the constant factor. For example, a subsequent version of the model, COINTOB, largely ignores this essential step as can be seen in their Fig. 1 (Brass et al., 2019). They write:

> *“[T]he threshold crossing is <u>mainly determined by</u> subthreshold neuronal noise […]”* (Brass et al., 2019, p. 256, underline added).

> *“A recent computational model […] suggested instead that random fluctuations of a motor readiness signal could be <u>sufficient</u> to explain the initiation of voluntary actions[…]”* (Ganos et al., 2015, p. 52, underline added).

> *“According to this model, the timing of the movement in the Libet experiment is <u>determined by</u> random threshold crossings in spontaneous fluctuations in neural activity. In particular, the model says that a decision when to move is <u>determined by random threshold crossings only</u> when it is not constrained by any evidence or reasons for action.”* (Schlosser, 2019, underline added).

Note that all these assertions would suggest that the imperative is 0 or close to 0, which is not how it is actually modelled. As shown above, it is possible – at least in principle – to provide a reasonable fit of the RP with zero imperative (Fig. 2). However, this does not provide a good fit for the decision times. It has been reported by the authors of the SDM that in order to obtain a good fit to *both* the waiting times *and* the shape of the RP within their model, the imperative is necessary (Guevara Erra et al., 2019). As we will see below, with the published SDM the imperative is not negligible, but an essential quantitative driving factor in threshold crossing.

If we look back to the perceptual decision-making case the roles are quite clear. When the sensory information level is high then the accumulator primarily integrates this evidence throughout the trial (the component determined by *I* in the model above). When the sensory information level is absent, *I* is set to 0 and the behavior is driven purely by the noise. In the SDM with the published parameters the situation is not like decision making without sensory information. *I* is not set to 0, so there is a constant driving input. However, as we have seen most of the interpretations in the literature focuses on the role of the noise fluctuations.

Some clarification is needed here, because the quantitative question of *how much* noise input versus constant input contribute to the crossing of the threshold can be dissected into three parts: First, as we have seen *both are necessary conditions* for reaching the threshold in a realistic time window. Second, given that the imperative is a constant factor, it is clear that within the SDM framework the trial-by-trial variation in response time and the shape of the waiting time distribution are explained by the random component that is entailed in the noise input. This is very similar to accumulator models for PDM, where the trial-wise differences in decision times are explained alone by the noise component. It is thus trivially clear that within the SDM framework the trial-by-trial variation in response time can *only* be explained by the stochastic component that is entailed in the noise input.

Third, we can assess the *quantitative contribution* of each of the two inputs, constant imperative and variable noise, to the crossing of the threshold. The question here is: How much overall input have the constant versus the noise components provided at the point in time when the threshold is crossed. It is essential here to take a close look at the model. In each time step the accumulator gets input from *both* the constant component (in the SDM the imperative) and the noise. Thus, movement towards the threshold is achieved by a combination of the continuous “push” of the constant and the variable (zero-mean) “rattle” of the random input. The combined and accumulated effect of these variables is additionally subject to a leak. One way to compare the contribution of the stochastic and the non-stochastic component to crossing of the threshold is to assess *how much of the distance the accumulator travels between zero and the threshold can be attributed to each component*. So we will sum up all the stepwise contributions of the noise component 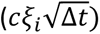 (i.e. the “rattle”) and also of the imperative component (*I*Δ*t*) (i.e. the “push”) separately up to the point where the accumulator crosses the threshold. This reflects the *input side* to the accumulator. Note that this is a stage prior to the leak (which in turn operates at the level of the combined accumulated signal, i.e. it affects the accumulated inputs of both noise and imperative).

A simple illustration of the net input of both sources is shown in Figure 5. It shows how much input to the accumulator has come from either the constant input (Fig. 5, black line), i.e. the continuous “push”, versus the noise input (Fig. 5, colored lines), i.e. the rattle. It is clear from Fig. 5 that the accumulated input from the constant is much higher than that of the noise. Note that the colored curves in Fig. 5 show the net accumulated input from (or integral of) the noise, not the noise itself. Note also that the time courses of the accumulated noise input wax and wane and are often below zero, as would be expected for zero mean noise.

Figure 6 shows a different perspective on this process, now viewed *backwards* from the time of threshold crossing. The histograms show the cumulative input between trial onset and threshold crossing separately for the constant imperative and for the noise (and separately for the two published variants of the SDM). For the first SDM version (Schurger et al., 2012) the imperative dominates the net input. It has a mean accumulated input of 0.7 (arbitrary units, averaged across 10000 trials). In contrast, the net contribution of the noise is on average 0 across all trials.

**Figure 6:**
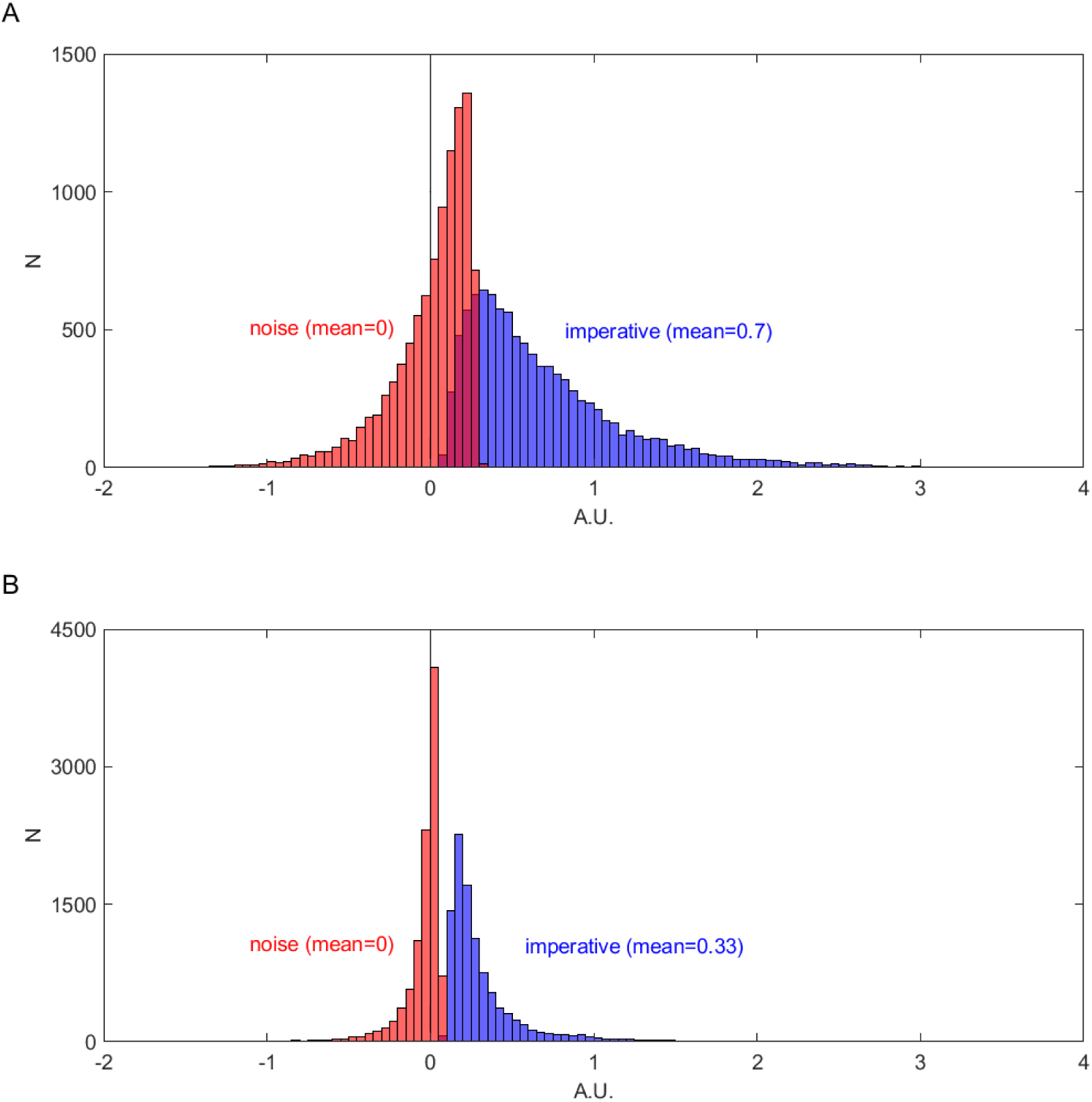
Total accumulated *absolute* contribution of imperative (dark gray, right histogram) and noise (light gray, left histogram) to the crossing of the threshold. The total contribution of imperative is always positive whereas the contribution of noise is in many trials even negative. The imperative dominates the overall input into the accumulator. **A** First version of the SDM (Schurger et al., 2012). **B** The finding is very similar for subsequent extended SDM (same scaling)(Schurger, 2018). Please note that within this model the trial-wise differences in decision times are a separate matter. They are explained by the noise and not by the imperative (see text).

How is that possible? An analogy might help here: Let’s assume a really strong person is pressing against a door to open it. They exert a strong force, but can’t quite get the door open. Then, a second, very weak person comes, does a very slight rattle on the door, whereupon the door jumps open. The combined force was sufficient to open the door, both are necessary. But the strong person makes the stronger quantitative contribution. It is similar for the SDM, where the constant push reflects the constant factor and the rattle is analogous to the noise. In the SDM, both factors are necessary, but quantitatively the constant factor makes the larger contribution (using the reported best fitting parameters from Schurger et al. 2012).

Figure 6 reveals that in many trials the net contribution of the noise is even negative. But how is it possible that the noise can have a negative net input if it is at the same time necessary to bring the accumulated signal cross the threshold? This happens when the noise is negative for long stretches of the trial and thus cancels out the positive input from the constant imperative. Then, towards the end of the trial, a small positive contribution can help the accumulated signal over the threshold (see also below for more details). Thus, in these trials the noise *prevented* the threshold from being crossed early (Fig. 6, A). For a second version of the SDM (Schurger, 2018, with pink noise as input) the findings are very similar. The imperative dominates the input with a mean of 0.33 (averaged across 10000 trials) and the noise contributes on average 0 (Fig. 6, B). The smaller values for the second model (Schurger, 2018) are due to a lower threshold compared to the first model (Schurger et al., 2012) (0.298 vs. 0.1256).

The relative strength of the noise and imperative obviously play a role here. To assess this, we repeated the simulation for different values of the constant factor *I* while keeping the noise scaling factor c and all other parameters constant (from Schurger et al. 2012) (see Fig. 7). At the time of reaching the threshold, the total input contributing to the accumulator is much higher from the constant imperative than from the accumulated noise for a wide range of values of I (Fig. 7, accumulated imperative = blue curve; accumulated noise = red curve). Noise only dominates the accumulated input for very small values of *I* close to 0 (i.e. where the red curve is above the blue curve). Trivially, when *I* = 0 the noise provides the only input and this hence has to be positive.

**Figure 7:**
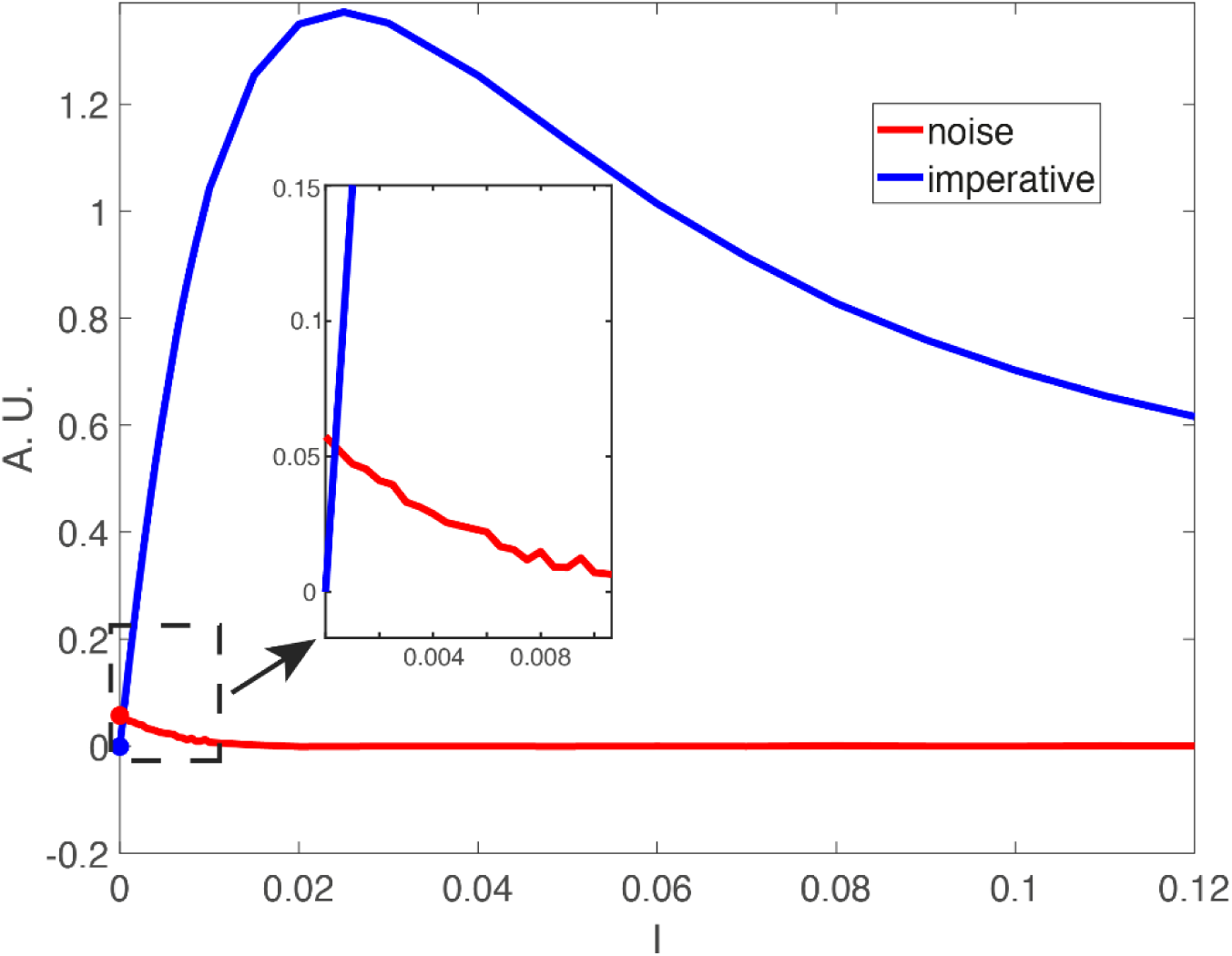
Total averaged accumulated *absolute* contribution of imperative (blue) and noise (red) to crossing the threshold for different values of *I* (imperative) in the model (all other parameters are constant and are taken from Schurger et al. 2012). For most values of *I* the accumulated total contribution of *I* exceeds the accumulated total contribution of the noise. Only for very small values of *I* close to 0 the contribution of noise exceeds the contribution of imperative. This is when the imperative is so small it hardly contributes to the drift towards the threshold (which could be achieved alternatively with noise with much higher variance). Note that the accumulated contribution of *I* rises first and then decreases again (with *I*). This is because the waiting times for low values of *I* are much longer and thus the amount of accumulated signal lost to the leak is higher. For higher values of *I* the waiting times are shorter, so the amount of accumulated signal lost to leak is smaller and thus the overall input is smaller. Please note that the contribution of the noise to the explanation of the trial-wise differences in decision times is a separate matter (see text). Also note that due to the leak (that affects the combined input of both sources) the total accumulated input has to be much higher than the threshold value.

In Figure 7 it is also clear that the average contribution of the noise across a wide range of parameters is close to 0, which is what would be expected for Gaussian noise with a mean of 0. As mentioned above, this does not exclude that the noise has a positive contribution in the brief time immediately preceding the threshold crossing.

The relative contribution of imperative versus noise to threshold crossing is also different for slow and fast trials. Thus, interestingly, the net noise contribution is negatively correlated with the waiting time. On short trials the noise contribution is positive. That is because the constant input alone will not have provided enough net input and thus short trials can only occur when the noise also has a positive input. In contrast, long trials can only come about if the constant positive input of the imperative is counteracted by a net negative contribution of the noise (i.e. otherwise the threshold would have been crossed earlier). Thus, the longer the trial the lower the accumulated net noise contribution for crossing the threshold.

We mentioned above that long trials occur when the noise counteracts the positive input from the imperative, and then only just before threshold crossing the noise provides a positive input. This raises the question how much the noise versus the constant input contribute to crossing the threshold in the last seconds before threshold crossing. This is also important because the SDM assumes that the readiness potential only emerges from the signals in the last 5 s of the time series. To address this, we will consider the quantitative contribution of noise versus constant in this final buildup period of the RP. We know that the noise has to have a positive contribution in that small time window. But does the noise input dominate over the constant input *at least in this final brief time window*?

To illustrate this, we show one selected simulated and exemplary trial of the SDM for illustrative purposes (Fig. 8). In the appendix we show that the following is also true for the averaged model RP. In Figure 8 we plot one simulated trial of the SDM with a long waiting time. For this trial we also show how much the noise (red) and imperative (blue) contributes from 0 at the trial start to the threshold crossing. A long waiting time can only be the result of a weak or even negative contribution of noise because the imperative is a constant positive input. As can be seen in Figure 8 the total input of noise (red line) is strongly negative going. Within this negative going noise there are small epochs in which the noise is slightly positive going. However, in most cases this is not enough positive contribution so that the SDM crosses the threshold. Only at the very end of the trial there is a small time window in which the noise can, together with the imperative, contribute enough, such that the threshold is crossed. Please note that in half of the trials the noise contribution is positive going (see Fig. 5). However, the general behavior is the same that the noise has the strongest positive contribution only very briefly before the threshold is crossed (for more details see the analyses in the Appendix).

**Figure 8:**
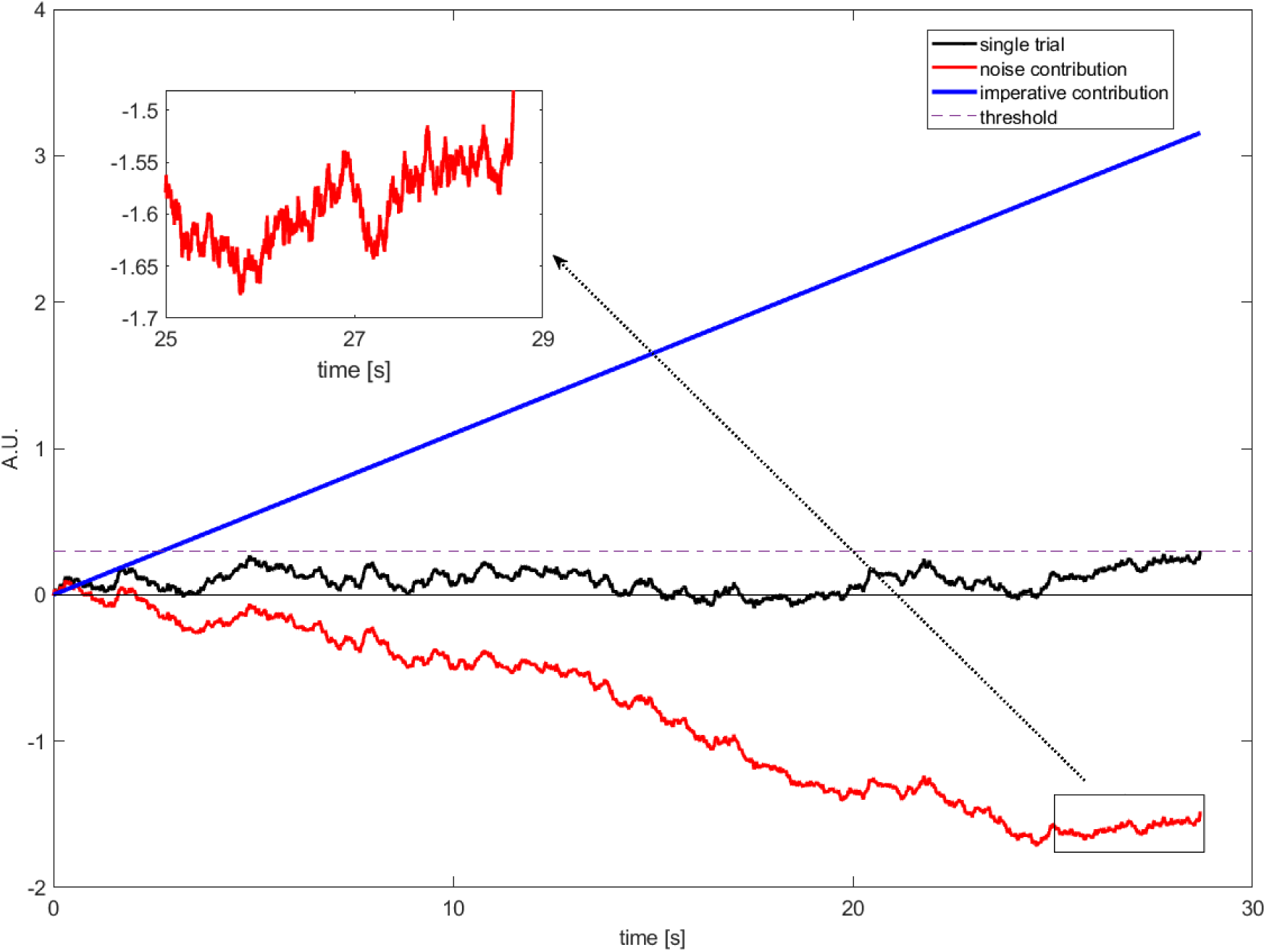
A single illustrative simulated trial of the SDM (black). The threshold is crossed relatively late at 28.7 s. The imperative contribution is constantly increasing and always “pushing” towards the threshold (blue). The noise contribution is negative going and counteracting the constant input, leading to a long waiting time for this simulated trial (red). The noise fluctuates (“rattles”) and there are small epochs during which the noise is pushing towards the threshold although on average it is negative going. Only at the very end of the simulated trial there is a short time window in which the noise has a contribution that is strong enough to push the SDM across the threshold together with the imperative.

However, one might consider our emphasis on the importance of the imperative a distraction. Why is this so important? There are important reasons to highlight the role of the imperative: in the model the imperative characterizes a fully deterministic component of the decision that is fixed once the trial has begun. The data by Murakami et al. (2014) also suggest that the information about when the decision is made is already there early in the trial. Furthermore, the LBA model where all the decisions are made at the beginning of the trial also predicts the RP and the distribution of waiting times (Brown & Heathcote, 2008). We will see below that these facts might change the interpretation about whether the decision is made early or late. As we will explore in the next section, there is a common view that the deterministic component is simply a preparatory stage, that brings the signal within reach of the threshold and then subsequently fluctuations take over. We will see that this is also not the case.

### Does the “imperative” (constant) term first bring the system into a dynamic range where random fluctuations take control?

As we have seen above there is another important aspect of the model, that noise- and imperative-related processes are interpreted as constituting separable and *sequential* stages (for examples see above). The idea there would be that the constant signal initially drives a “*stochastic exponential transition period”* (Schurger et al., 2012, p. E2906) that first brings the accumulated signal into an operating range, and subsequently the fluctuations determine when the signal crosses the threshold (Schurger et al., 2012, p. E2906). Here are a few examples of this point:

> *“After a stochastic exponential transition period […], the leaky accumulator generates noisy trajectories whose threshold crossings determine movement times.”* (Schurger et al., 2012, p. E2906);

> *“In our model this solution amounts to simply shifting premotor activation up closer to the threshold for initiation of the instructed movement and waiting for a random threshold-crossing even*t.” (Schurger et al., 2012, p. E2905);

> *“According to their stochastic decision model, the decision process, given Libet’s instructions, amounts to simply shifting premotor activation up closer to the threshold for initiation of the movement and waiting for a random threshold-crossing fluctuation in RP.”* (Bayne & Pacherie, 2015, p. 224).

Considering these statements, we should expect two effects: First, the accumulator is moved closer to the threshold without any (or only few) decisions being made. Second, from this plateau the system waits for a random threshold-crossing event.

Let us consider the time point five seconds into the trial where the accumulator has on average reached around 90% of its asymptote (see Fig. 9, middle). One might assume that hardly any decisions have been made by this point, but quite the opposite is the case. In 52.7 % of the trials the threshold is crossed and a decision is made earlier than 5 s. For the trials with long waiting times it can even be observed that the signal fluctuates strongly and sometimes even reaches negative values after it was first closer to the threshold (see the orange and yellow curves on the right of Fig. 9, bottom). Therefore, the “move signal closer to threshold” process can even happen multiple times in slow trials. Thus, the verbal description and interpretation in the quotation above of noise and imperative does not capture the model behavior appropriately. During the entire time course of a trial both imperative and noise contribute to the current state of the accumulator. It’s a concert of the two, both contributing to the process, with the imperative exerting a larger overall quantitative contribution towards reaching the threshold, and the noise explaining trial-wise differences in decision time. Please note, that above arguments also hold if time points earlier than 5 s are considered as reaching a lenient interpretation of a plateau. We observe many early decisions and the SDM signals fluctuate strongly.

**Figure 9:**
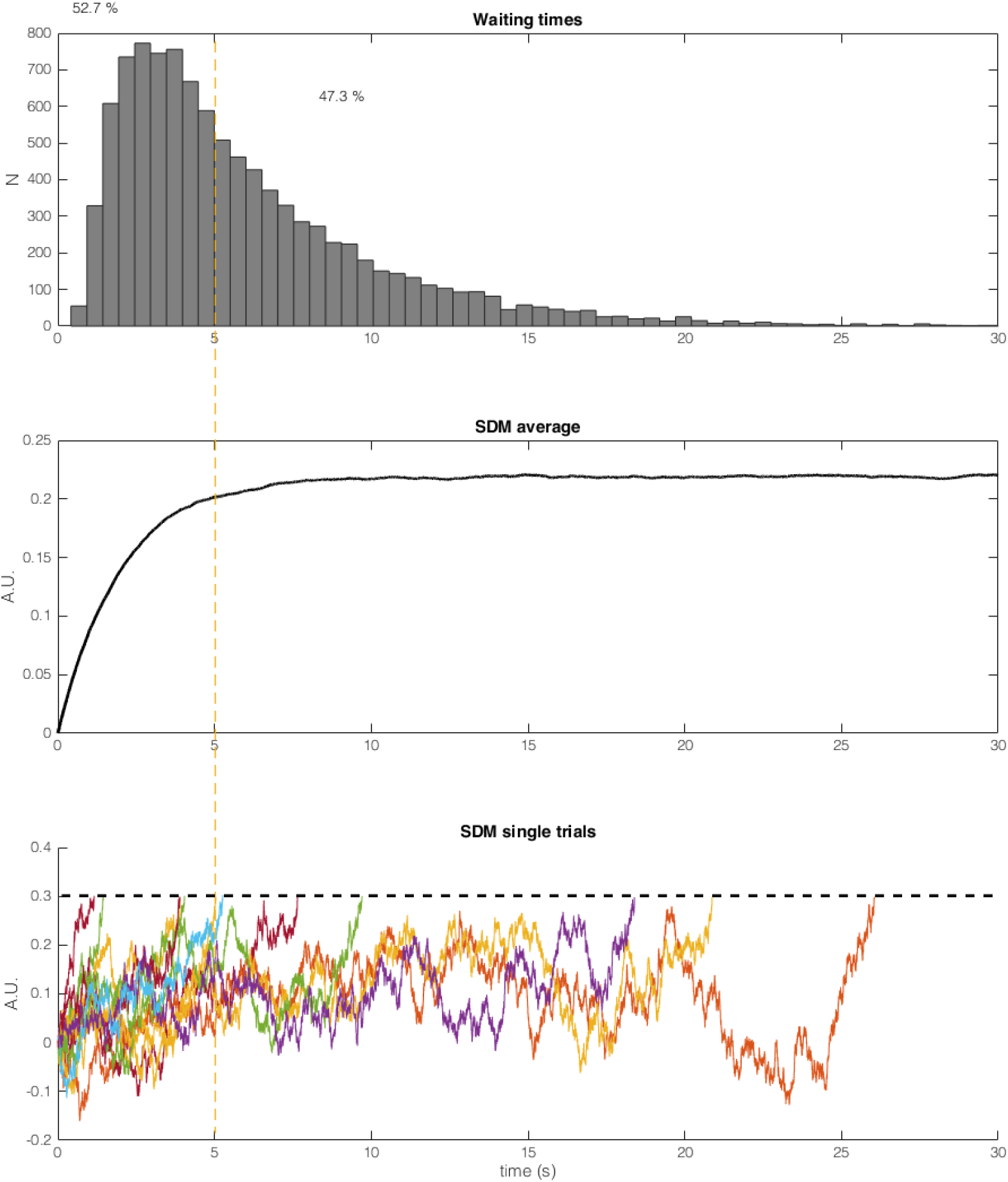
Result of 10000 trials of the SDM calculated with the parameters reported in Schurger et al. (2012). **Top:** Histogram of waiting times. **Middle:** Average SDM output of 10000 trials. The average SDM converges after around 5 s. In the averaged signal, the noise across trials cancels out and the result is similar to a SDM with only imperative and no noise as input (Fig. 1). **Bottom:** 10 sample trials of the SDM (truncated after crossing the threshold, dashed line). More than half of the trials (52.7 %, left side of orange line) cross the threshold before 5 s. Trials with long waiting times don’t stay close to the asymptote but fluctuate strongly. Neither trials with a waiting time faster than 5 s nor trials with a very slow waiting time shows the proposed behavior that the signal is moved closer to the threshold and that then some noise causes a threshold crossing. The imperative and noise both continuously influence the SDM signal, while the imperative is only driving the signal up, noise is driving the signal up and down.

Please note, in the extended version of the SDM with pink instead of white noise as input (Schurger, 2018) the logic of the two stages is even more problematic. There, the constant imperative can drive the SDM across the threshold alone, i.e. without the noise, and thus two sequential stages are not necessary anymore. Here is the proof: we can calculate *x*_*i*_ at the asymptote, i.e. when Δ*x*_*i*_ is zero. We also consider the case without noise.

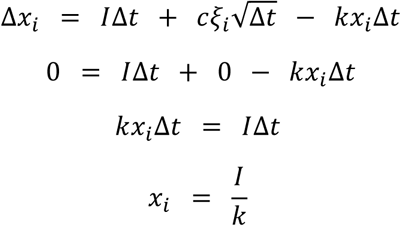

With the reported parameters (I = 0.1, k=0.6 and threshold = 0.1256) the SDM would converge to 0.1/0.6 = 0.167 based on imperative alone and without noise. The threshold in this model is at 0.1256 so the threshold would be crossed without any noise in the model. Taken together, as mentioned above, it is not accurate to think of the SDM as a two-stage model, with phase 1 being “climb closer to threshold (but don’t cross it yet)”, and phase 2 being “OK, now you may cross the threshold at any time”.

### Is any “evidence” involved in the model?

In perceptual decision making, “evidence” refers to one variable having information *about* another, such as a perceptual representation having evidence about an external stimulus. This is the reason it is called “evidence” and not simply “a signal in MT” or “bias”. In contrast, in the SDM the imperative describes an intrinsic signal that is not evidence, but a signal that is necessary for the total accumulated signal to cross the threshold in behaviorally realistic times (Guevara Erra et al., 2019). In line with this, the authors of an animal study on endogenous movement decisions that they consider to reflect an accumulation process (Murakami et al., 2014), say that their task involves “*no evidence per se”* (p. 1580).

There seems to be some confusion in the literature about whether one of the signals (imperative *or* noise) might reflect evidence in the SDM after all. Already an early review paper interpreted the original study as

> *“showing that bounded-integration processes, which involve the accumulation of <u>noisy evidence</u> until a decision threshold is reached, offer a coherent and plausible explanation for the apparent pre-movement build-up of neuronal activity.”* (Schurger et al., 2016, p. 77, underline added).

Brass, Fürstenberg & Mele (2019) interpret the original paper on the SDM as a solution that:

> *“treat[s] stochastic noise in the motor system as <u>evidence</u> for the accumulation process”* (Brass et al., 2019, p. 256, underline added).

They then continue:

> *“In contrast to perceptual decision making, however, the accumulation of evidence [in the SDM] is not based on perceptual information but on <u>internal information</u> and stochastic neural activity.”* (p. 259, underline added).

And then:

> *“These models assume that decision time in the Libet task is based on a process of accumulation of evidence to a threshold, just like in other decision-tasks. Because the decision is not based on perceptual or other external evidence, this <u>accumulation of evidence might operate primarily on stochastic neural fluctuations in the motor system</u>”* (p. 257, underline added).

And finally:

> *“This means that the RP and the LRP do not reflect a ballistic process that necessarily leads to action but rather a <u>gathering of evidence</u>.”* (p. 259, underline added).

Thus, it appears that some authors consider that the noise plays a role of evidence, and that there is some additional signal involved, here termed “internal information”. Also in other papers there is some ambiguity as to the respective roles of the variable factor (i.e. the noise fluctuations) and the constant factor (evidence/imperative):

> *“Schurger et al. propose that the motor system constantly undergoes random fluctuations of RPs and that this random premotor activity is used as a substitute for actual evidence.”* (Pacherie, 2014, p. 36).

Of course it is possible to go beyond the original formulation of the SDM and re-consider the imperative signal as having some computational function dedicated to representing decision-relevant internal states (such as motivation or impulsivity). One possibility could be that there is a single overall mechanism, but with two different types of input, one being sensory evidence and the other being imperative.

When considering one variable having evidence about another one would want it to fulfill some additional requirements. For example, the evidence should be able to “stand in” as a proxy of what it is representing (Shea & Shea, 2018, p, 15-16, 143). To illustrate this, we may turn to Brass et al. (2019), who note that it could indeed be sensible to assume that some latent internal signals could influence the buildup of the imperative when no external information is available. In such cases, the level of imperative could be somewhat constrained by these causally influencing factors, but we would not necessarily see the imperative as having a function of reliably tracking such variables and serving as a stand-in (i.e. “evidence”) for those. Not every causal influence can be considered as evidence. Please further note that if the imperative indeed played a role of collecting evidence it would also have needed to receive much more attention as an integral part of the movement decision, and not be largely ignored as we have seen above.

### What does it mean for the decision to be early or late?

As mentioned above, a classical interpretation of the readiness potential is that it reflects a *post*-decisional stage of processing after an unconscious decision to act has been made (Schurger et al., 2012). First the brain makes an *early* decision, then a process (of which the RP is an indicator) is triggered that prepares the movement. Some consider this to be counterintuitive because of the long temporal delay of many hundreds of milliseconds between the brain’s unconscious decision and the time when a participant consciously believes to be “making the decision now” (Libet et al., 1983). An important reason for the interest in the stochastic decision model is that it seems to remove this counterintuitive time delay. It implies instead that the decision to act occurs *late*, that is when the accumulated signal crosses the threshold for action (Brass et al., 2019; Schurger et al., 2012), which is much closer to the subjective time of decision. Everything happening before that is pre-decisional activity in the brain (some of it stochastic, according to the SDM). In this view the readiness potential originates at a *pre*-decisional stage and is an artefact of averaging stochastic signals aligned to the time of a threshold crossing.

Just as a reminder, “decision” here can mean two things: (a) the participant’s conscious experience of making a decision, and (b) some (potentially unconscious) brain event, quasi a “neural decision”^1^, that somehow sets the brain on the track for executing the movement (these two events may or may not coincide in time). Because the discussion of the SDM has focused on the threshold crossing, which is a property of the neural system, we will focus on neural decisions and set the problem of subjective decisions aside. And in order to avoid a too extended general discussion of the role of randomness and (in)determinism in biological systems, we will base our discussion on properties of the computational models. The question we want to discuss here is whether accumulator models always imply late decisions. We would like to suggest that the interpretation of the *nature of the fluctuations* in the SDM is vital when interpreting the decision as early or late. Please note that this is not specific to the SDM though, but it holds for any stochastic accumulator model.

In the papers on the SDM the fluctuations are described as “internal physiological noise” or “random fluctuations” (Schurger et al., 2012, p. E2905 and E2904). This could mean different things, so an important distinction is required. **(a)** Either this could mean that they are “*objectively random”* in the sense of the randomness being an irreducible part of the world that would not be predictable in more detail however much we learn about the world. An extreme version of this were if fluctuations were guided by quantum processes (analogous to e.g. radioactive decay)^2^. Due to their indeterminism such fluctuations could not be predicted. Whatever one might find out about the states of the world, there is an intrinsic indeterminism that remains. **(b)** On the other hand the randomness could refer to “*epistemic randomness”*, where a signal would *appear* to have random properties, but this would reflect the fact that some properties of the process are not known (e.g. due to insufficient data or to not understanding the algorithm, say as in the case of a deterministic random number generator). The process could also be unpredictable despite being deterministic (as for example in deterministic chaos). The randomness might also constitute a mixture of **(a)** and **(b)**, so we might uncover more latent determinants as research progresses, but there still remains an irreducible remainder.

Now let us see whether the interpretation of the nature of the processes leading up to the threshold influences the time at which we would consider the decision to happen. In order to cleanse our thinking of the a-priori assumptions we have with accumulator models, let us consider a physical analogy, a Rube Goldberg-style simple chain reaction model of a ball rolling down a slope towards a row of dominos (Fig. 10). The dominos fall over one by one and then ring a bell. In each trial the ball is set off with a slightly different speed and we measure the time until the bell is rung. Note that when the ball knocks over the first domino stone a nonlinearity is reached, such that a different process is triggered. If the ball for some reason were prevented from reaching that stage, say it were to mysteriously move uphill again, then the chain reaction in the dominos would not be triggered. Thus, knocking over the first domino is similar to passing a nonlinear threshold (analogous to *β*).

**Figure 10:**
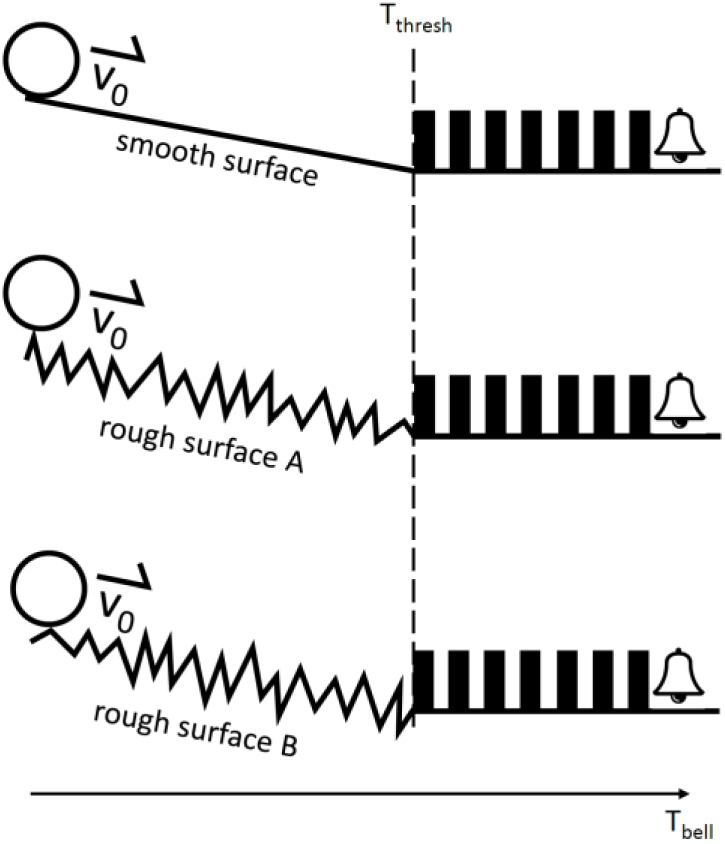
Simple causal chains inspired by Rube Goldberg to illustrate the models. **Top:** A ball is set off on an even slope with variable speed v_0_ that is determined at the beginning of the trial from a Gaussian random distribution. The slope is chosen to exactly balance the friction and keep the ball at a constant speed without acceleration or deceleration. At the end of the slope at time T_thresh_ the ball triggers a chain of dominos that runs deterministically through and then sounds a bell at time T_bell_. Note that if the ball (for some reason) were to be stopped from reaching the dominos they would not be triggered, so the passing of the threshold can be considered a nonlinear event. When would the decision be made when the ball reaches the bell? There is only one free variable: the speed with which the ball is initially set off. So is the decision made at the beginning of the trial? **Middle:** Now the same causal chain is set off but with an added rough surface. Now the time that is needed to pass the rough slope influences the time until the bell is rung. But is this process deterministic? That depends on the nature of the rough surface. One type of surface would potentially result in slightly different bumps on every trial and thus add some irreducible variability to the run time. A different type of surface might still potentially yield reproducible run times. **Bottom:** Now, let’s stick with the reproducible version and exchange the rough slope on a trial-by-trial fashion, each time with a different reproducible slope. Let’s assume we don’t know which slope has been picked on a given trial. When is the decision about the run time made? Given that following the release of the ball (analogous to the beginning of a trial) everything is fully deterministic, one plausible interpretation would be to say that the decision is made at the beginning of the trial, despite appearing to depend on random processes on the way. Even though the model only reproduces some properties of the accumulator models (for example there is no leak) it can help form our intuitions about which events might count as decision points. Without an in-depth understanding of the nature of the noise and without an ability to measure it, it is impossible to say whether a process is deterministic or not. And whether the decision is early or late will presumably depend on the answer to this question.

We will consider four different cases: **(1)** In the first case (Fig. 10, top) a ball rolls down a constant *smooth* slope, knocks over the first domino, which triggers a predictable chain of events until the last domino rings the bell. Everything in this model is perfectly determined at the beginning of the trial when the ball is set off with a certain speed. **(2)** In the second case (Fig. 10, middle) everything is the same, with the exception that the surface is now rough. Let’s assume that the rough surface causes the ball to bump back and forth on the way, thus adding a random component to the time it takes to pass the ramp. Let’s further assume that the effect of these bumps is indeterministic in principle, so that however precise we can measure the ball and the surface there would still remain some degree of indeterminacy. **(3)** Now let’s consider a third case, similar to (2) but now the effect of the bumps is perfectly deterministic. When the ball is let go with a certain speed and the same rough surface the run time is always the same. **(4)** Let’s also consider a variant on (3). Now in every trial we use a different rough slope (Fig. 10, middle and bottom). Each of these slopes is perfectly deterministic, but we don’t know which one is chosen on a given trial. Case 1 approximately symbolizes the LBA, case 2, 3 and 4 represent variants of the SDM.

Cases 1 and 3 are clearly deterministic. Case 2 is indeterministic. And case 4 is concealed determinism that appears as indeterminism because a latent variable (which slope is chosen) is not known. The rolling down the slope is an analogy to the drift phase and the domino stones are an analogy for the motor execution stage triggered after passing the threshold. If all the relevant causal factors are established (as in cases 1, 3 and 4), isn’t the outcome of the decision then pre-determined and it can thus be considered already made? It would be interesting to conceptually clarify this issue for interpreting the implication of stochastic processes in general.

### Summary and outlook

The aim of this paper was to assess the level of evidence for the SDM and to address certain confusions that have arisen in the literature. First, we highlighted that there is no direct evidence based on neuronal-level measurements for the role of stochastic fluctuations in the RP and movement initiation and that the analyses are based on macroscopic signals averaged across many trials. This does not rule out the SDM as a model, but it clarifies what kind of data will be required to definitively validate the model. Second, we found that the purported evidence for the SDM from animal studies is limited and may not favor the SDM over other models. Third, we showed that a quasi-deterministic model where the parameters are fixed at the beginning of the trial (the LBA) makes very similar predictions to the SDM, including for the interrupted version of the Libet task, and fits well with a population of neurons found in an invasive monkey study. Fourth, we found that the literature has tended to ignore the deterministic component of the SDM, the imperative signal, that accounts for an important part of signal input towards the threshold. Both the stochastic fluctuations and the imperative are necessary for reaching the threshold in realistic time periods with the published parameters of the SDM. Fifth, there is a confusion regarding the link between perceptual decision making and spontaneous movements. We have argued that the SDM is not just a special case of perceptual decision making, but without the evidence. Although mathematically identical to a leaky stochastic accumulator used to model perceptual decision making, the SDM does not incorporate any “evidence” per se (as assumed in some secondary sources, see above). In the context of the SDM, the “evidence” in perceptual decision-making is replaced by a different constant factor, an imperative to move given by the demand characteristics of the task. To avoid confusion, we recommend using the term “imperative” in the context of the SDM rather than “evidence”. Sixth, we remind the reader that a key aim of the SDM, to provide a pre-decisional account of the RP, cannot be fully addressed by the model because this hinges critically on the nature of the noise fluctuations as being “objectively random” versus “epistemically random”. This of course is true of any scientific model that incorporates randomness, and may be very difficult to decide empirically, but at least the case is far from closed.

There are some other questions that need to be addressed: **Where is the ramp?** Is there any insight into the neural mechanisms underlying the large initial imperative-based ramp at the beginning of the trial where the averaged accumulator signal moves closer to the threshold? If this process involves the same neural mechanism as the readiness potential (i.e. it reflects the signal in the accumulator), then one would expect to see a very stereotypical ramp at the beginning of the trial. In fact, invasive recordings from human single neurons have revealed a slow gradual ramping up of the signal prior to the time of decision (Fried et al., 2011).

**Could the ramp not be the buildup of the intention?** The fact that the imperative signal plays such an important role in bringing about the decision could point to a re-interpretation of the components of the SDM model. One way would be to interpret the imperative signal as the largely deterministic and gradual buildup of the intention to move (Schurger et al., 2016) and the smaller effect of randomness some form of intrinsic variability. Would this mean that the decision is made early or late? Random variability from trial to trial is observed in just about any task (from threshold perception to motor performance) without this randomness necessarily being considered the most relevant property of the process.

It could also be useful to extend the scope from thinking about spontaneous movements in general to **what happens specifically in spontaneous movement *experiments***. These lab experiments impose constraints that are not present in real-world free-ranging actions. For example, there is an explicit or implicit affordance to move within a reasonable time-frame (e.g. to not wait too long) and at the same time avoid being predictable or rhythmic, which Schurger et al. (2012) refer to as the demand characteristics of the task and incorporate in the model as the imperative (or drift) term in the SDM. Already the earliest paper on readiness potentials stated: “*The participant was required to perform the movement not rhythmically, but in irregular intervals”* (Kornhuber & Deecke, 1965, p. 1, our translation). In the study by Schurger et al. (2012), the instructions are to “*[…] try not to decide or plan in advance when to press the button, but to make the event as spontaneous and capricious as possible.”* (Schurger et al., 2012, p. E2911). What if the participant is thinking: “Oh dear, am I spontaneous or capricious enough?”, and if so what would they do? In the classic Libet study the participants are required to “[…] *let the urge to act appear on its own at any time without any preplanning or concentration on when to act”* (Libet et al., 1983, p. 625). One might wonder what participants were thinking if they didn’t experience such a mysterious “urge”. Would they have just waited for the whole duration of the experiment and then finished the experiment by saying: “*Sorry, but I never felt an urge to move”*? As has been pointed out previously (Schurger et al. 2012, Supporting Information) the key point here is that the preparation of these movements might have involved a vast array of cognitive processes, conscious or unconscious. These could include (among others) mental time keeping and time-based prospective memory (McDaniels & Einstein, 2000), inhibition of behavioral impulses to move immediately (Noorani & Carpenter, 2017), or generation of random behavior sequences (Nickerson, 2002). Sticking with the latter, obviously, in order to be random and capricious one could in principle use a random time interval generator based on the accumulation of fluctuations. But why generate or use a long series of random numbers, if all you have to do is generate one single random number at the beginning of the trial (as e.g. in the LBA)?

In a recent review paper the authors summarize the evidence for the SDM:

> *“Why should there be such a long and highly variable lag, of up to one second or more, between the decision to initiate movement and movement onset? Why has the RP not proven to be a very reliable real-time predictor of movement onset? And why are subjective reports of the (conscious) decision time so late relative to the supposed ‘onset’ of the RP? These puzzling questions are not at all puzzling from a late-decision perspective, so the onus should be on proponents of the early-decision account to explain why the more parsimonious late-decision view is false.”* (Schurger et al., 2021, p. 5).

These questions might have simple answers. For example, the delay after the brain’s early decision to move until the movement occurs could simply reflect a delay that is decided upon already at the beginning of the trial and that is introduced to comply with the instructions of the task, similar to the LBA. Please note that this entails a commitment to participants being mistaken about not having prepared the movements ahead of the time of their subjective decision. That would also predict why the RP onset is not a reliable predictor of the movement onset. Of course, this assertion further entails a commitment to the assertion that subjects are patently and reliably wrong in their subjective reports of not pre-planning, or that the decision is made pre-consciously, as Libet (1983) asserted. Furthermore, the LBA model is arguably no less parsimonious than the SDM because it involves fewer variables (i.e. it does without the unmeasured and thus hypothetical fluctuation time series). We would like to clarify that we do not want to argue that the verdict is already in for an early decision model (as in the LBA), or that the SDM can be ruled out based on the evidence. At the current state of evidence, the debate between early-decision and late-decision accounts is still not settled. We point out that the empirical support for the model is currently not yet definitive, and that also several key conceptual issues still need clarification. We hope we have provided some of that clarification here.

## Appendix

Here we will calculate the average quantitative contribution of noise versus constant in this final buildup period of the RP. We know that the noise will typically have a positive contribution in that small time window. But does the noise input dominate over the constant input *at least in this final brief time window*?

To address this, we will consider the model RP resulting from back averaging from the threshold crossing event in 1000 trials (see above main text). We separate the model RP into its 3 additive components (imperative, noise and leak) based on the core equation of the SDM, repeated here for convenience:

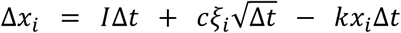

Please note that we can use the equation also for the average model RP and not just single trials. From the given model RP we can calculate average Δ*x*_*i*_. We also know *x*_*i*_ (which is the model RP at time step i) and thus can calculate the average leak (*kx*_*i*_Δ*t*). Then we can calculate the average noise contribution because the imperative is constant. By repeating this calculation for every time step of the model RP we can separate the different contributions. We separated the model RP starting from 3 different time points (-2, -1 and -0.5 s). Finally, we cumulated the 3 different components from the starting points (-2, -1 and -0.5 s) to 0 s (when the threshold is crossed) (see Fig. A1, top row). Please note that we extend this analysis and look at the contribution of the average leak as well because the model RP is the sum of all the three components.

**Figure A1:**
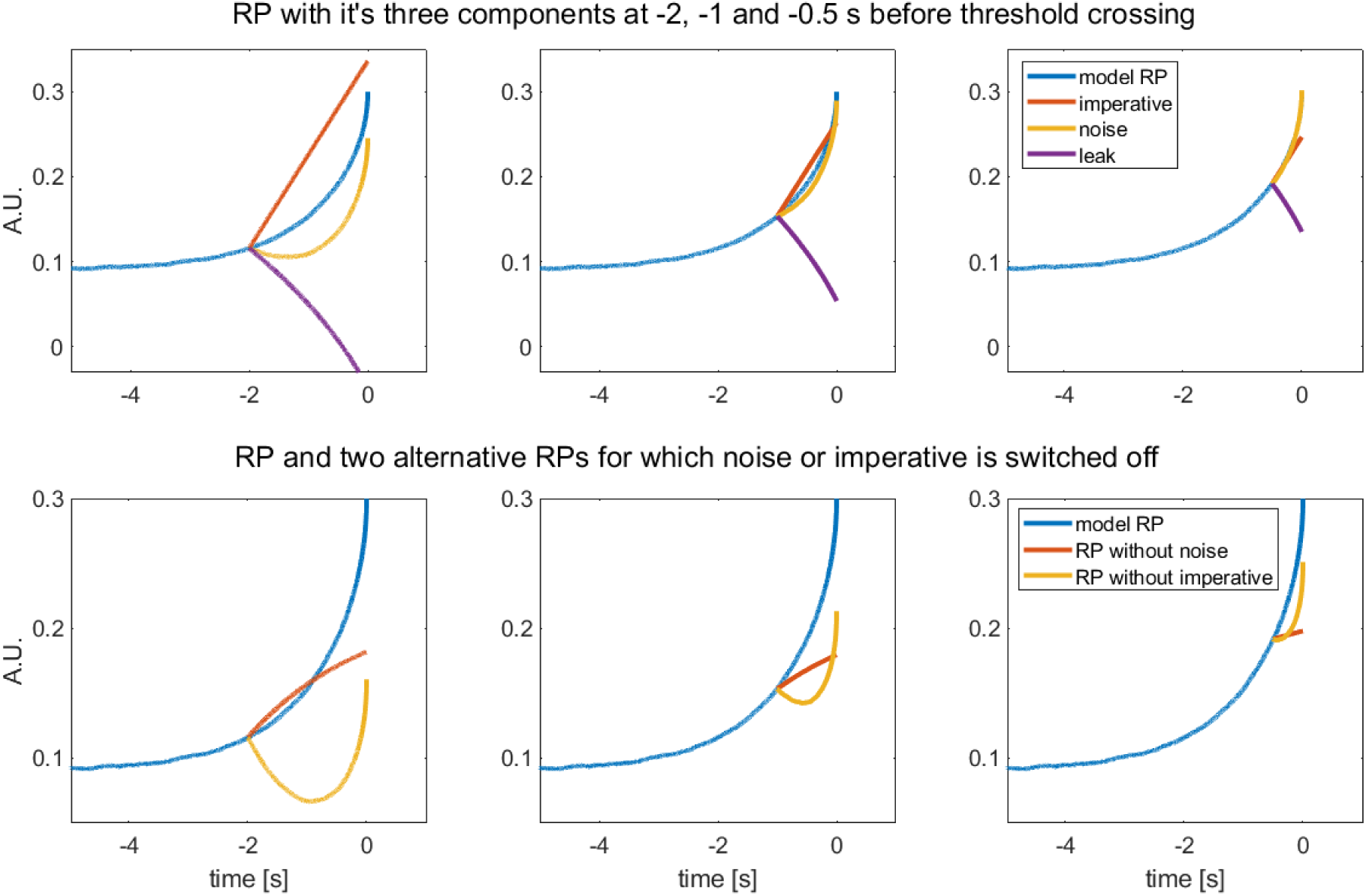
Top: Model RP (blue) and the separated contribution of imperative (red), noise (yellow) and leak (purple) into the model RP for 3 time windows (-2 to 0, -1 to 0 and -0.5 to 0 s from left to right). The contribution of imperative:leak for the 3 time windows is 0.22:0.13, 0.11:0.14 and 0.06:0.11. The later and shorter the time window the higher the relative contribution of noise (compared to imperative) with a higher contribution of noise compared to imperative for the shorter time windows. Bottom: RP (blue) and alternative versions of the RP for which the noise (red) or imperative (yellow) input was set to 0 at -2, -1 or -0.5 s (from left to right) before crossing the threshold. None of the model RPs would have crossed the threshold of 0.298. Even for the shortest time window of 500 ms in which the average noise contributes roughly twice as much compared to the imperative, the constant input is needed to cross the threshold.

Please note that this is a biased analysis regarding the average noise contribution. Just before crossing the threshold the noise must have a positive contribution. Thus, although the noise contribution across the whole trial might be negative in a single simulated trial (see above) in a small time window before crossing the threshold the noise contribution has to be positive. Consequently, it is not surprising due to the selection of the time windows before averaging, that the average noise has a positive contribution. However, it can be seen that for the longest of the three time windows (-2 to 0 s) the imperative still dominates the input (Fig. A1 top left). Only for the shortest time window (-0.5 to 0 s) the average noise clearly dominates the input over the imperative (0.11 vs. 0.06 respectively) (Fig. A1, top right).

In a next analysis we calculated two alternative model RPs for which we set either the imperative or the noise to 0 at three different time points (-2, -1 and -0.5 s). We used the average noise estimates from the first analysis and recalculated the leak according to the equation of the model. With this analysis we simulated how the model RP would change if one of the 2 main input sources would have been switched off in order to illustrate their role for crossing the threshold and for the shape of the model RP (Fig. A1, bottom row). The leak in the model is relatively strong (see Fig. A1, top row purple). Therefore, switching off the constant input of the imperative leads to a strong decrease of the alternative model RP at first (Fig. A1, bottow left, yellow). This further shows how small the input of the average noise is, as it cannot compensate for the pull of the average leak down to zero, until roughly 500 ms before the threshold is crossed when the alternative model RP without imperative still increases. It is also very important to note that even for the shortest time window (Fig. A1, right) in which the average noise contributes more than the imperative, the average noise is not strong enough to push the accumulator over the threshold (0.298) without the constant push of the imperative. In other words, switching the imperative off 500 ms before the decision would not have generated a decision (with the reported parameters). Please note that the noise plotted in Figure A1 is the average noise and therefore it appears smooth without the typical “rattle”.

## Acknowledgements

This work was funded by the Excellence Initiative of the German Federal Ministry of Education (Excellence Cluster Science of intelligence), the BMBF (through the Max Planck School of Cognition), the DFG (GRK 2386 “Extrospection”), and a joint grant by the John Templeton Foundation and the Fetzer Institute. We would especially like to thank Aaron Schurger who acted very constructively as an adversarial partner. His invaluable collegial input has helped us make very important clarifications in the manuscript. We would also like to thank Matthias Schultze-Kraft, Thomas Goschke, Marcel Brass and Michael Pauen for valuable discussions on the topic.

## Footnotes

1 We use the term “decision” here without further discussion, but consider a clarification of what could constitute a decision without making reference to the subjective experience a major challenge in this field. From a neural level of resolution, brain processes can be described as trajectories in a high-dimensional state-space. It is unclear what would constitute a decision along such a trajectory. If one considers phase transitions or bifurcations as decisions, as criteria then these would be ubiquitous.

2 At least in indeterministic interpretations of quantum mechanics.

